# A nuclease-driven mechanism of post-replicative ssDNA gap suppression

**DOI:** 10.1101/2025.10.22.683685

**Authors:** LG Bennett, V Thanendran, D Harrison, CM Jones, E Antonopoulou, J Rajan, EG Vernon, A Gamble, R Kraehenbuehl, E Hartsuiker, CJ Staples

**Affiliations:** North West Cancer Research Institute, North Wales Medical School, Bangor, Gwynedd, Wales, UK LL57 2UW

## Abstract

During replication stress, failure to resolve post-replicative ssDNA gaps generated by PRIMPOL-mediated replication repriming is linked with chemosensitivity, and in all models reported to date the nuclease MRE11 has been implicated as a gap-promoting factor. We have dissected ssDNA gap dynamics following nucleoside analogue-mediated nascent strand termination and report a novel mechanism via which loss of the MRE11 negative regulator MRN-Interacting Protein (MRNIP) leads to MRE11 exonuclease-dependent suppression of post-replicative ssDNA gaps. This process is driven by UBC13 and the B-family TLS polymerase REV3L, suggesting that in the absence of MRNIP, dysregulated MRE11 activity at chain termination sites licenses gap filling via template switching. In the absence of PRIMPOL-dependent repriming, MRNIP loss leads to paradoxical SMARCAL1, MRE11 and primosome-dependent ssDNA gaps, suggesting that nucleolytic digestion of reversed forks generates DNA intermediates that platform primase redundancy in the context of chain termination. We also highlight novel site-specific roles for the anti-resection factor 53BP1 in enabling primase redundancy via suppression of the nuclease EXO1, and in limiting the EXO1-dependent processing of post-replicative ssDNA gaps. Finally, we demonstrate that CDK-dependent MRNIP phosphorylation is required for MRNIP functionality in the regulation of ssDNA gaps and sensitivity to chain terminators. This work represents the first report of nuclease-driven post-replicative gap filling, illuminates an additional level of versatility in the replication stress response, and expands our understanding of the context-dependent links between nuclease regulation and chemoresistance.

## Introduction

Elegant and complex mechanisms elicit DNA damage repair and replication stress tolerance. Among these, DNA replication repriming by the DNA primase-polymerase PRIMPOL facilitates continued DNA synthesis in response to template strand impediments^1–5,29^. PRIMPOL-mediated repriming leads to the generation of ssDNA discontinuities in nascent strands, which must be subsequently filled by post-replicative gap filling mechanisms such as Template Switching (TS) or Trans-Lesion Synthesis (TLS). Gap-filling pathway choice is cell cycle phase-dependent, proceeding either via UBC13-triggered PCNA poly-ubiquitination and S-phase TS, or by RAD18-driven PCNA mono-ubiquitination and G2-phase TLS^6,7^. A range of factors have been implicated in gap suppression, including the canonical tumour suppressors BRCA1 and BRCA2^7–12^. Loss of either BRCA1 or BRCA2 function leads to increased ssDNA gap prevalence linked to chemosensitivity, driven by the exonuclease activity of MRE11 (3’-5’) and EXO1 (5’-3’)^13^. We recently demonstrated that MRN-interacting protein (MRNIP) limits MRE11 exonuclease activity against dsDNA substrates *in vitro* and identified a role for MRE11-Interacting Protein (MRNIP) in the limitation of MRE11-dependent post-replicative gaps induced by cross-linking agents in cells deficient in replication fork remodelling^14^. Gap-filling in each of these models is mediated by the processive B-family polymerase REV3L, the catalytic component of the TLS Pol-ζ complex^7^.

Chain-terminating nucleoside analogues (CTNAs) such as Gemcitabine and Cytarabine are widely employed in the treatment of pancreatic cancer, bladder cancer and several leukaemia subtypes. CTNAs mediate cytotoxicity at least in part via inhibition of DNA synthesis, following their direct incorporation into the nascent DNA strand. Recent evidence suggests that the exonuclease function of the DNA repair factor MRE11 contributes to Gemcitabine removal from genomic DNA^15^, although the dynamics of DNA replication in response to chain termination remains relatively understudied.

Here, we demonstrate the unique functionality of MRNIP in the context of CTNA-induced nascent strand synthetic blockade. We previously reported that MRNIP loss leads to sensitivity to multiple replication stress-inducing agents, including Poly-(ADP-ribose) Polymerase (PARP) inhibitors and the Ribonucleotide Reductase (RNR) inhibitor hydroxyurea (HU)^14,19^. In contrast, MRNIP KO cells are highly *resistant* to Gemcitabine and the obligate chain terminator Cytarabine but are sensitive to other NAs such as Fludarabine and Clofarabine. MRNIP loss results in suppression of Gemcitabine and Cytarabine-induced ssDNA gaps via a mechanism driven by UBC13, REV3L and the exonuclease activity of MRE11. Recent evidence indicates that the anti-resection factor 53BP1 can influence gap-filling pathway choice in *BRCA1*-mutant cells^37^, and our findings are consistent with the hypothesis that in addition to Gemcitabine removal, a minimal threshold of MRE11-dependent resection is required to stimulate post-replicative gap filling by TS.

We also observed paradoxical 53BP1, MRE11 and SMARCAL1-dependent DNA gaps in PRIMPOL-depleted MRNIP KO cells, which were also suppressed by the concomitant depletion of primosome subunits. These observations lead us to posit that defective repriming at chain termination sites can be resolved via fork reversal and the subsequent end-resection of nascent DNA ends to generate a template for redundant repriming by the remaining functional primase. In further support of this hypothesis, we identify context-specific roles of the anti-resection factor 53BP1 in the control of EXO1-dependent end resection at PRIMPOL-dependent and PRIMPOL-independent gaps. Our data suggests that bidirectional loss of end resection control upon dual deficiency in MRNIP and 53BP1 prohibits post-replicative gap filling, and furthermore, that primase redundancy is contingent upon template maintenance facilitated by 53BP1-mediated EXO1 repression.

Our work also sheds light on the regulatory mechanisms that support MRNIP function. We demonstrate that MRNIP Ser217 phosphorylation is essential for MRNIP functionality in the regulation of chemosensitivity and post-replicative gap prevalence in response to Gemcitabine. Control of Cyclin-Dependent Kinase 1 (CDK1) activity has been linked to Gemcitabine sensitivity. For example, small-molecule inhibition of the CDK1 negative regulators WEE1 and PKMYT1 sensitise cancer cells to Gemcitabine^16–18^. These findings suggest a wider role for CDK activity in the regulation of post-replicative gap processing. In summary, we have identified a novel chemoresistance mechanism via which loss of a nuclease regulatory factor facilitates post-replicative ssDNA gap-filling and chemoresistance.

## Materials and Methods

### Cell culture and CRISPR-Cas9 cell line generation

WT and MRNIP KO HeLa and HCT116 cells were maintained as adherent monolayers in Dulbecco’s Modified Eagle’s medium (DMEM) with 10% Foetal Bovine Serum at 37°C in an atmosphere of 5% CO_2_. Stable MRNIP KO CRISPR clones were generated via transfection of HeLa cells with Santa Cruz Biotechnology CRISPR-Cas9 gRNA and HDR plasmids (sc-412131-KO-2 and sc-412131-HDR-2). The following day, the cells were reseeded into 10-cm plates in the presence of puromycin (2 μg/ml). After a further 10 days, individual clones were picked, subcultured, and analyzed by PCR from gDNA, qRT-PCR, and Western blotting to confirm deletion of MRNIP. The characterisation of the clones used in this study has been previously reported^19^.

MRNIP KO cells stably expressing PRIMPOL were generated via co-transfection of pDEST-V5 WT or CH mutant primase-dead PRIMPOL and pOG44 recombinase constructs (a kind gift from the Petermann laboratory, Birmingham, UK)^20^, followed by selection in 150 μg/ml hygromycin.

### RNAi transfections

Cells were transfected with 10-50 nM siRNA according to cell type using Lipofectamine RNAiMAX (Invitrogen) according to the manufacturer’s instructions. Cells were subsequently further treated, lysed or fixed for analysis at least 48 hrs post-transfection unless otherwise indicated.

### siRNA sequences

Control: 5′-UAAGGCUAUGAAGAGAUAC-3′

PRIMPOL: 5′-GAGGAAAGCUGGACAUCGA-3′

PRIMPOL 5′-UTR GGAGAUGGACAACGUAUUU-3′

UBC13 5′-GGCUAUAUGCCAUGAAUAA-3′

RAD18 5′-CCAAGAAACAAGCGUAAUA-3′

REV3L 5′-GGAGUUCUCUGCUGAGUUA-3′

53BP1 5′-GAAGGACGGAGUACUAAUA-3′

SMARCAL1: 5′-GCUUUGACCUUCUUAGCAA-3′

MRNIP: 5′-GUUAGGAGGGACAGGGUUC-3′

PRIM1: 5′-GCAAAGGAAUCAAUCAUCU-3′

POLA1: 5′-GAUGGUAAAGCACGCAAUA-3′

POLA2: 5′-UGAGAGAUGUGCACCAUGA-3′

EXO1: 5′-GCACGUAAUUCAAGUGAUG-3′

### Cell lysis and Western blotting

For the preparation of whole-cell extracts, cells were solubilized in lysis buffer [25 mM Tris-HCl (pH 7.4), 150 mM NaCl, 1% Triton X-100, 1 mM dithiothreitol (DTT), and 1 mM MgCl_2_] supplemented with Benzonase (50 U/ml) (Novagen) and cOmplete protease inhibitors and PhosSTOP phosphatase inhibitors (Roche). Lysates were clarified by centrifugation at 16,000*g* for 15 min at 4°C. Gel electrophoresis was performed using 4-12% NuPAGE gels (Invitrogen). Briefly, samples were resolved in NuPAGE MES running buffer and transferred to polyvinylidene difluoride (PVDF) membranes, which were then probed for the protein of interest using antibodies diluted in phosphate-buffered saline (PBS)–0.1% Tween 20 (Sigma-Aldrich) containing 5% Marvel or 5% BSA as appropriate. Blots were imaged on a Bio-Rad ChemiDoc machine.

### Immunofluorescence

Cells were grown on glass coverslips in 24-well trays, transfected/treated as indicated, fixed with 3% buffered paraformaldehyde for 10 min at room temperature, and permeabilized in PBS containing 0.5% Triton X-100 for 5 min at room temperature. Cells were blocked in 3% BSA and incubated with primary antibody overnight in the cold room and subsequent incubation with Alexa Fluor 488– or Alexa Fluor 594– conjugated goat anti-rabbit or anti-mouse immunoglobulin G fluorescent secondary antibodies (Invitrogen, 1:1000). Antibody dilutions and washes after incubations were performed in PBS. DNA was counterstained with 4′,6-diamidino-2-phenylindole (DAPI, 1 μg/ml), and coverslips were mounted cell-side down in Shandon Immu-Mount medium (Thermo Fisher Scientific). Fluorescence microscopy was performed on a Zeiss LSM710 confocal microscope at ×40 or ×63 magnification. Images were captured and analysed using Zen software (Zeiss).

### Native IdU Assay

Cells were grown on glass coverslips in 24-well trays, transfected with relevant siRNAs, and after 24 hrs cells were incubated with 25 μM IdU for 24 hrs. Cells were then treated with genotoxins and/or inhibitors as required, and fixed in 3% buffered paraformaldehyde in PBS for 10 min at room temperature. Indirect immunofluorescence was performed using an anti-mouse BrdU antibody (BD, clone 3D4, 1:250) that cross-reacts with IdU, an Alexa anti-mouse 488 secondary antibody (Invitrogen, 1:1000) and DAPI counterstaining.

### Clonogenic survival assays

Cells were plated onto 6-well trays and transfected with siRNAs as required. Twenty-four hours later, 1000 cells were replated onto 10 cm culture dishes and treated with genotoxic agents as required after a further 24 hrs. Colonies were incubated for 12 days then fixed and stained using methylene blue/methanol. Colonies were counted manually, and results normalised to untreated controls.

### DNA fibre assay

Cells were plated and transfected appropriately, then were pulse-labelled with 25 μM CldU (Sigma-Aldrich) and 250 μM IdU (Sigma-Aldrich) as indicated in combination with replication stress-inducing agents and/or inhibitors as required. Cells were then permeabilized with CSK buffer (100 mM NaCl, 10 mM MOPS, 2 mM MgCl_2_, 300 mM sucrose and 0.5% Triton X) for 5 min, then washed in S1 buffer (30 mM sodium acetate, 10 mM zinc acetate, 5% glycerol and 50 mM NaCl), then treated with 20 U/mL S1 nuclease (Thermo) in S1 buffer for 30 min. Cells were then harvested in PBS. Nuclei were then pelleted and resuspended in PBS. Two point five microlitres of suspended nuclei were mixed with 7.5 μl of lysis buffer (200 mM Tris pH 7.4, 25 mM EDTA, and 0.5% SDS) on a clean, dry slide (Thermo Fisher Scientific). After 7 min, slides were tilted at 30°, air-dried, then fixed in cold methanol/acetic acid (3:1). DNA fibres were denatured using 2.5 M HCl for 75 min then washed with 1× PBS before blocking in 3% Bovine Serum Albumin (BSA)/PBS containing 0.2% Tween 20 for 1 hour. CldU- and IdU-labelled tracts were incubated with two anti-BrdU (5-bromo-2′-deoxyuridine) antibodies, one of which is specific for CldU (Abcam) and the other for IdU (BD). Slides were then washed and incubated with goat anti-mouse/rat Alexa Fluor 488 and Alexa Fluor 594 (Invitrogen). DNA fibres (150 per condition) were visualized on a Zeiss LSM710 confocal microscope, and images were collected using Zen software and then analyzed with ImageJ.

### List of antibodies

The following antibodies were used: Western blotting: PRIMPOL (Proteintech: 29824-1-AP, 1:1000), RAD18 (Cell Signalling, D2B8, 1:1000), UBC13 Antibody (Santa Cruz, F-10, 1:1000), GAPDH Antibody (Santa Cruz, G-9, 1:2000), MRNIP Antibody (Santa Cruz, H-11, 1:500), phospho-S217 (in-house, generated by Eurogentec, 1:1000), Tubulin (Abcam, ab4074), SMARCAL1 (Santa Cruz Biotechnology, sc-376377), γH2AX (Cell Signaling Technology, 9718), PRIMPOL (gift from Mendez laboratory, 1:1000). Immunofluorescence: 53BP1 (Abcam, ab21083, 1:2000), IdU (BD, 3D4, 1:250) and γH2AX (Upstate, 1:1000).

### Quantitative RT-PCR

RNA was extracted using RNeasy Plus kits (Qiagen, Cat No. 74134), and 1 μg RNA was reverse transcribed using a Superscript^TM^ III First-strand Synthesis kit (Thermo-Fisher Cat. No. 11752050) according to the manufacturer’s instructions. PCR reactions were set up in triplicate and reactions run on the Bio-Rad CFX system using the following validated Quantitect primers (GeneGlobe IDs: GAPDH (QT00079247) and REV3L (QT00044653), respectively) and GoTaq qPCR Master Mix (Promega Cat. No. A6001). Cycling parameters were as follows: 95°C for 2 minutes, then 40 cycles of melting and extension (95°C for 15 sec and 60°C for 1 minute). Data was collected and analysed on CFX Manager software (Bio-Rad). Triplicate Ct values for targets (REV3L) were normalised to Ct values for GAPDH.

## Results

### MRNIP loss leads to CTNA resistance and DSG suppression

We previously observed that in the absence of fork reversal, loss of MRNIP leads to an increased prevalence of PRIMPOL and MRE11-dependent post-replicative ssDNA gaps and PRIMPOL-dependent chemosensitivity ^14^. While triaging the response of MRNIP KO cells to various genotoxic agents, we performed cell viability assays in WT and MRNIP KO HeLa and HCT116 cells following treatment with the CTNA Gemcitabine. Unexpectedly, MRNIP KO cells of both backgrounds were Gemcitabine-resistant (Fig 1A and Supplementary Fig 1A). Drug resistance was not an off-target effect of the CRISPR protocol, since stable expression of FLAG-tagged MRNIP in a MRNIP KO background restored chemosensitivity to levels observed in WT cells (Figure 1A). Gemcitabine mediates chemosensitivity via both chain termination and RNR inhibition (in the diphosphate form), and we previously demonstrated that MRNIP promotes resistance to the RNR inhibitor Hydroxyurea (HU)^19^. To begin to clarify the mechanism underpinning the observed resistance to Gemcitabine, we performed viability assays following treatment with Cytarabine, which is an obligate chain terminator^21^. Notably, MRNIP KO cells were resistant to high doses of Cytarabine (Fig 1B), in contrast to the gradual loss of viability observed with increasing Gemcitabine concentration. This is consistent with dNTP pool perturbation at higher doses of Gemcitabine. In further support of this hypothesis, MRNIP KO cells were sensitive rather than resistant to the purine analogues Fludarabine and Clofarabine (Fig 1C and Supplementary Fig 1B). Finally, we tested the viability of MRNIP KO cells following treatment with the topoisomerase I poison Camptothecin (CPT). MRNIP KO cells were sensitive to CPT, and resistance was restored by stable expression of FLAG-MRNIP in a MRNIP KO background (Supplementary Figure 1C).

**Figure 1:**
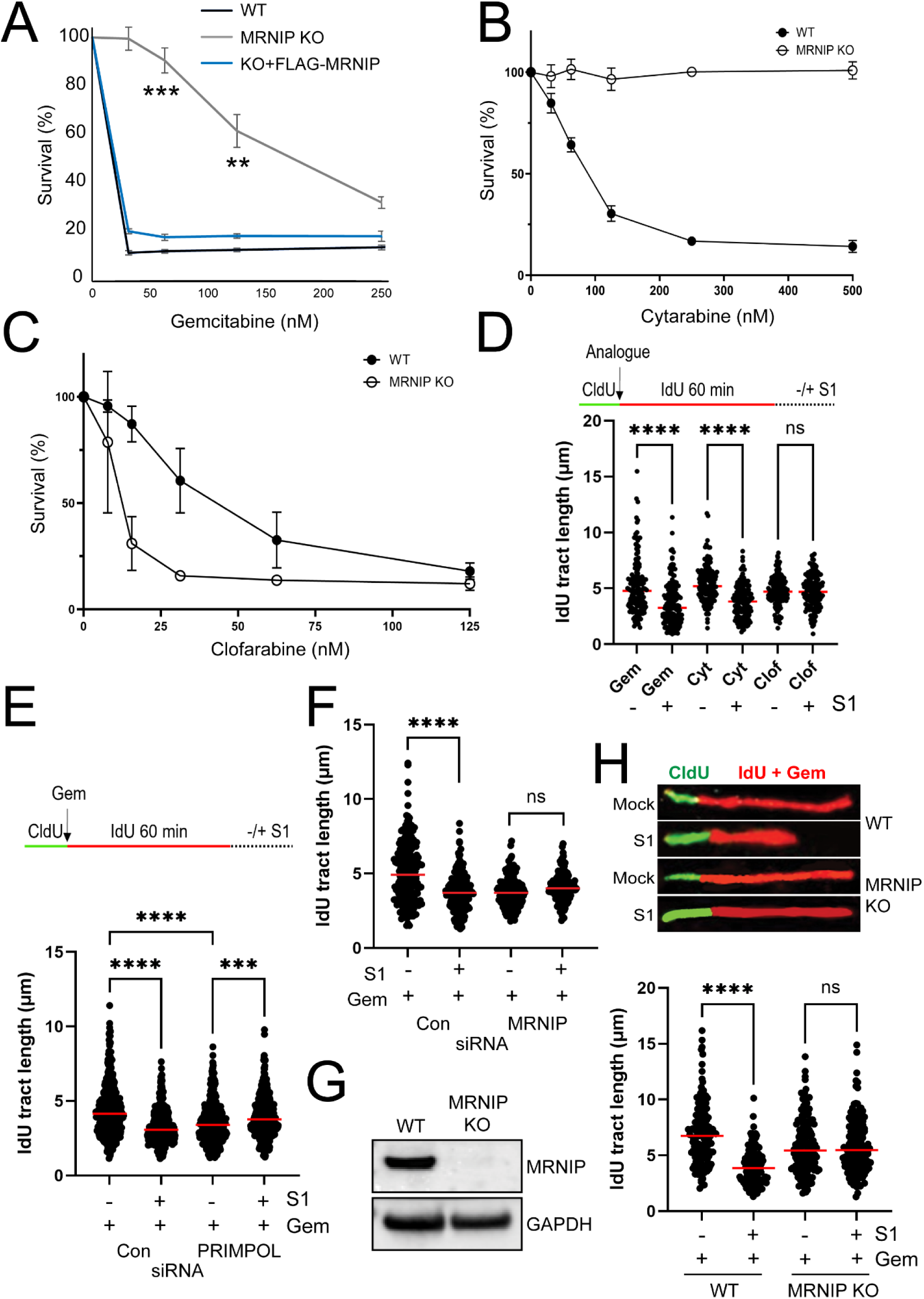
MRNIP loss leads to CTNA resistance and ssDNA gap suppression. **A-C:** WT and MRNIP KO HeLa cells and MRNIP KO HeLa cells stably expressing FLAG-MRNIP (A) were treated with the indicated concentrations of Gemcitabine, Cytarabine or Clofarabine. After 72 hrs, cell viability was estimated by MTT assay, and data normalised to untreated controls. **D:** Nascent DNA in WT HeLa cells was CldU-labelled for 20 mins, followed by IdU labelling for 1 hr in the presence of 100 nM Gemcitabine, Clofarabine or Cytarabine. Single-stranded DNA was then digested via S1 nuclease treatment, and DNA fibre spreading was performed. IdU tract length in the presence and absence of S1 was used as an indicator of the presence of post-replicative ssDNA gaps. **E:** WT HeLa cells were transfected with a control siRNA, or an siRNA targeting PRIMPOL. After 48 hrs, cells were labelled and treated as in (D). **F:** WT HeLa cells were transfected with a control siRNA or an siRNA targeting MRNIP and after 48 hrs, cells were labelled and treated as in (D). **G:** MRNIP status was validated in WT and MRNIP KO HeLa cells by Western blotting. **H:** WT and MRNIP KO cells were labelled with halogenated nucleoside analogues in the presence of 100 nM Gemcitabine and treated and analysed as in (D). All experiments were performed independently at least three times and A, B, E, G and H were independently confirmed by multiple laboratory members. Statistical analysis was conducted by one-way ANOVA. ns=not significant, Statistical analysis was conducted by one-way ANOVA. ns=not significant, *p<0.05, **p<0.01, ***p<0.001, ****p<0.0001

There is a paucity of published data on the generation and processing of post-replicative DNA gaps in cells treated with nascent strand terminators. Given the established role of MRNIP in the restraint of post-replicative gaps in other models, we set out to investigate a potential role for MRNIP in the regulation of gaps induced by nascent strand terminators. Initially, we assessed the presence of DNA gaps using S1 nuclease-coupled DNA fibre assays. The S1 nuclease cuts ssDNA, and as such IdU tract shortening following S1 nuclease treatment is indicative of the presence of post-replicative DNA gaps. To account for any potential effects on overall replication progression, we included mock-treated samples processed under identical conditions. Treatment of WT HeLa cells with Gemcitabine or Cytarabine resulted in the generation of DNA gaps as evidenced by IdU tract shortening post-S1 treatment (Fig 1D). In contrast, treatment with Clofarabine did not lead to DNA gaps, suggesting that CTNA-induced DNA gaps occur consequent to chain termination (Fig 1D). Depletion of PRIMPOL via RNAi demonstrated that the DNA gaps observed in Gemcitabine-treated HeLa cells are formed consequent to PRIMPOL-dependent repriming (Fig 1E and Supplementary Fig 1D). We went on to assess the prevalence of DNA gaps in MRNIP-deficient cells following Gemcitabine treatment. Notably, Gemcitabine-induced DNA gaps were completely suppressed in both MRNIP-depleted HeLa cells RNAi (Fig 1F) and MRNIP KO HeLa cells (Fig 1G and H).

### Gap suppression in MRNIP KO cells is dependent on the MRE11 exonuclease

The exonuclease activity of MRE11 has been implicated in Gemcitabine removal^15^. Given that Gemcitabine removal is a prerequisite for nascent chain extension, the detection of post-replicative DNA gaps (Fig 1D-G) in Gemcitabine-treated HeLa cells suggests that end processing and gap-filling following repriming at CTNA-blocked ends is relatively inefficient in WT cells and is at least delayed beyond the 1 hr timescale of our S1 nuclease-linked fibre experiments. Our previous studies demonstrate that MRNIP represses MRE11 exonuclease activity against dsDNA substrates *in vitro*, and that MRNIP loss results in increased nascent DNA degradation at reversed forks and PRIMPOL-dependent post-replicative ssDNA gaps in response to HU and PARP inhibition, respectively^14,19^. We therefore hypothesised that MRNIP regulates the MRE11-dependent processing of nascent DNA at sites of Gemcitabine incorporation. To test this, we performed S1-linked DNA fibre assays in Gemcitabine-treated MRNIP KO HeLa cells following pre-treatment with the MRE11 exonuclease inhibitor Mirin. In WT cells, exonuclease inhibition had minimal effect on Gemcitabine-induced ssDNA gaps. Remarkably, pre-treatment with Mirin led to restoration of Gemcitabine-induced ssDNA gaps in MRNIP KO cells (Fig 2A). To further assess these phenotypes, we performed native IdU immunofluorescence assays. Under these conditions, the IdU-specific antibody is capable of antigen recognition only in exposed regions of ssDNA. We observed increased ssDNA signal in WT cells following Gemcitabine treatment, which was suppressed in MRNIP KO cells and was unaffected by pre-treatment with the MRE11 endonuclease inhibitor PFM03 (Supplementary Fig 2A). Consistent with our S1-linked fibre assay findings, ssDNA levels in MRNIP KO cells were restored by pre-treatment with either PFM39 or Mirin, which are both specific inhibitors of the MRE11 exonuclease (Fig 2B)^22,23^. Subsequent experiments confirmed that PFM39 partially re-sensitised MRNIP KO cells to Gemcitabine (Fig 2C). Similar experiments using Mirin were confounded by toxicity associated with long-term treatment. Furthermore, MRE11 exonuclease inhibition with either Mirin or PFM39 led to a significant and selective increase in the proportion of MRNIP KO cells positive for the DNA damage markers 53BP1 and γH2AX (Fig 2D and E). Together, this work suggests a paradoxical role for MRE11 exonuclease activity in DSG suppression in Gemcitabine-treated MRNIP KO cells.

**Figure 2:**
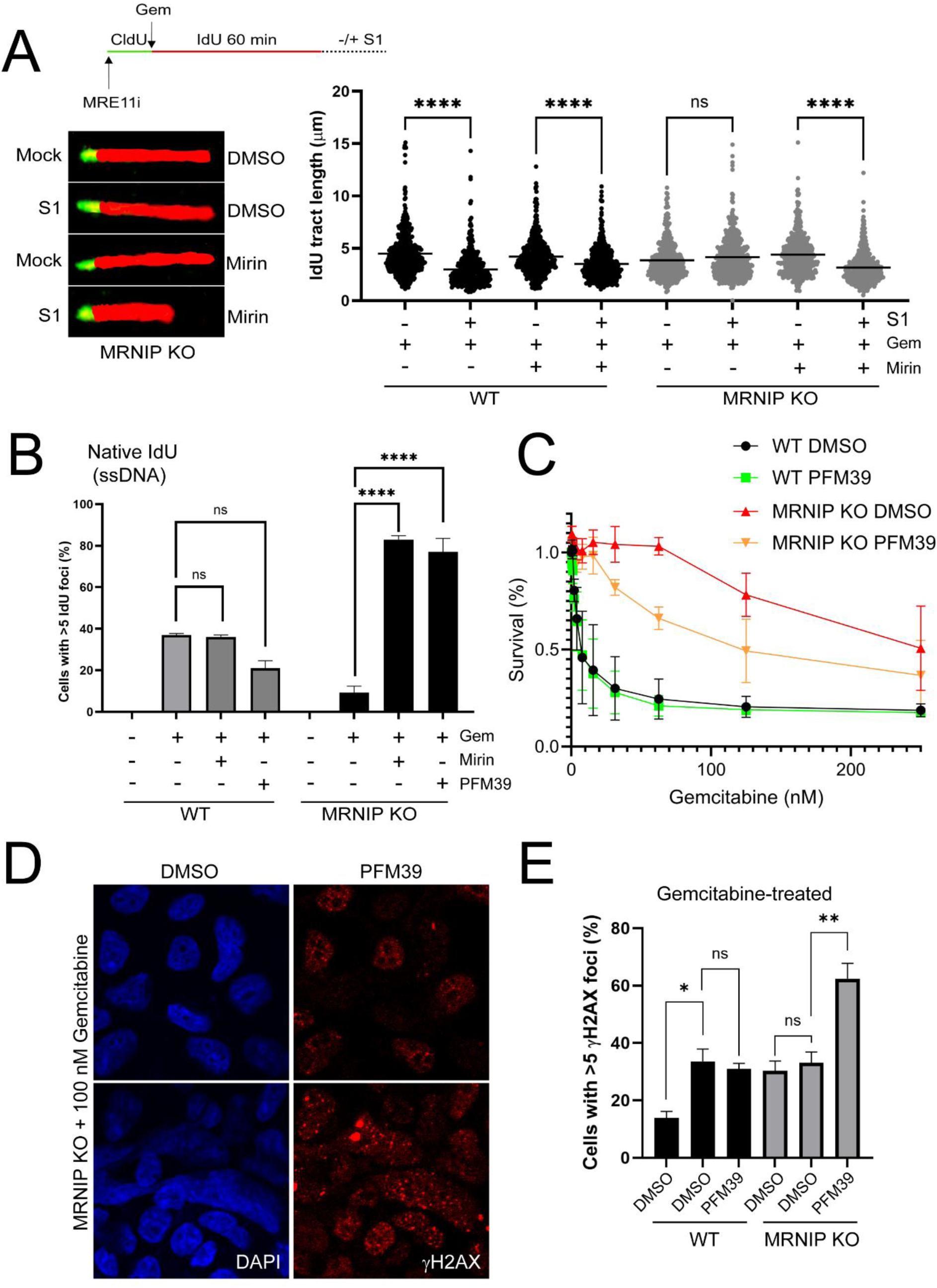
Gap suppression in the absence of MRNIP is MRE11-dependent. **A:** Nascent DNA in WT and MRNIP KO HeLa cells was CldU-labelled for 20 mins, followed by IdU labelling for 1 hr in the presence of 100 nM Gemcitabine and either DMSO or 25 μM Mirin. Single-stranded DNA was then digested via S1 nuclease treatment, and DNA fibre spreading was performed. IdU tract length in the presence and absence of S1 was used as an indicator of the presence of post-replicative ssDNA gaps. **B**: WT and MRNIP KO HeLa cells were labelled with IdU for 16 hrs, followed by 16 hrs treatment with vehicle control or 100 nM Gemcitabine in the presence of DMSO or 25 μM PFM39. Cells were then fixed and stained under native conditions with a IdU cross-reactive BrdU antibody, and counterstained with DAPI. Cells were assessed by confocal microscopy, and the proportion of cells with >5 nuclear IdU foci was determined. **C**: WT and MRNIP KO cells were treated with the indicated concentrations of Gemcitabine in the presence of either DMSO or 25 μM PFM39. After 72 hrs, cell viability was estimated via MTT assay. **D and E**: WT and MRNIP KO cells were treated with 100 nM Gemcitabine in the presence of either DMSO or 25 μM PFM39. After 24 hrs, cells were fixed and stained with a γH2AX antibody and counterstained with DAPI. Cells were assessed by confocal microscopy, and the proportion of cells with >5 nuclear foci was determined. Representative images are displayed in (D). All experiments were performed independently at least three times and A, D and E were independently confirmed by multiple laboratory members. Statistical analysis was conducted by one-way ANOVA. ns=not significant, *p<0.05,**p<0.01, ****p<0.0001

### Gap suppression in MRNIP KO cells is driven by template switching-mediated gap filling by POL-ζ

Our data is consistent with the hypothesis that MRNIP limits MRE11 nuclease activity at chain termination sites, and that MRNIP loss triggers enhanced MRE11-dependent resection, which subsequently facilitates post-replicative DSG filling. To test the involvement of replication repriming and post-replicative gap filling mechanisms in DSG suppression in this model, we performed DNA fibre and native IdU assays in Gemcitabine-treated MRNIP KO HeLa cells following depletion of the primase-polymerase PRIMPOL, the TS factor UBC13, or REV3L (the catalytic subunit of Pol-ζ). Native IdU analysis revealed that Gemcitabine treatment led to detectable ssDNA in WT HeLa cells (Fig 3A). As before, we could not detect nuclear ssDNA foci in Gemcitabine-treated MRNIP KO cells. However, following PRIMPOL, UBC13 or REV3L depletion, the majority of MRNIP KO cells displayed nuclear ssDNA foci (Fig 3A). In contrast, depletion of the TLS factor RAD18 failed to restore ssDNA gaps, consistent with previous reports that RAD18 promotes TLS in G2-phase, while TS fills post-replicative gaps behind the fork in S-phase. These observations were further supported by the restoration of DNA gaps as detected by S1-linked DNA fibre assays performed in Gemcitabine or Cytarabine-treated MRNIP KO HeLa cells depleted of UBC13, PRIMPOL or REV3L (Fig 3B and Supplementary Fig 2B), and in native IdU assays in MRNIP KO HCT116 cells (Supplementary Fig 2C).

**Figure 3:**
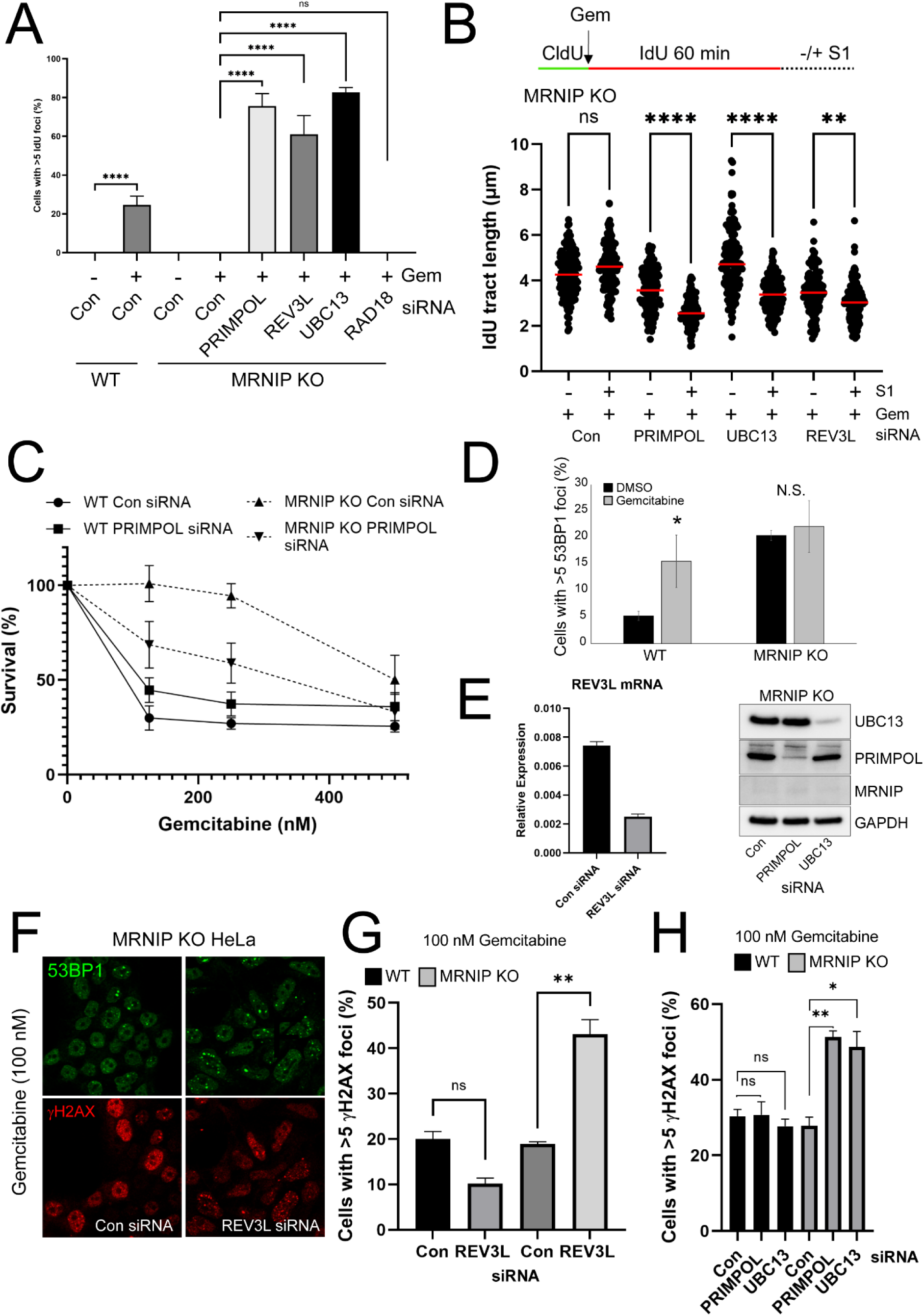
DNA gaps and DNA damage suppression in Gemcitabine-treated MRNIP KOs is driven by repriming and template switching. A: **WT and MRNIP** KO HeLa cells were transfected with a non-targeting control siRNA, or siRNAs targeting UBC13, PRIMPOL, RAD18 or REV3L. After 48 hrs cells were labelled with IdU for a further 16 hrs, followed by 16 hrs treatment with vehicle control or 100 nM Gemcitabine. Cells were then fixed and stained under native conditions with a IdU cross-reactive BrdU antibody, and counterstained with DAPI. Cells were assessed by confocal microscopy, and the proportion of cells with >5 nuclear IdU foci was determined. **B**: MRNIP KO cells were transfected with the indicated siRNAs. After 48 hrs, nascent DNA in WT and MRNIP KO HeLa cells was CldU-labelled for 20 mins, followed by IdU labelling for 1 hr in the presence of 100 nM Gemcitabine. Single-stranded DNA was then digested via S1 nuclease treatment, and DNA fibre spreading was performed. IdU tract length in the presence and absence of S1 was used as an indicator of the presence of post-replicative ssDNA gaps. **C:** WT and MRNIP KO HeLa cells were transfected with a control siRNA or an siRNA targeting PRIMPOL. After 48 hrs, cells were replated and treated with the indicated concentrations of Gemcitabine and assessed 72 hrs later via MTT assay. **D**: WT and MRNIP KO HeLa cells were treated with 100 nM Gemcitabine for 16 hrs prior to fixation and staining for the DNA break marker 53BP1. Cells with >5 nuclear 53BP1 foci were scored as positive. **E:** Validation of REV3L, PRIMPOL and UBC13 depletion via western blotting and qRT-PCR. **F-H:** WT and MRNIP KO cells were transfected with a control siRNA or siRNAs targeting either PRIMPOL, UBC13 or REV3L. After 48 hrs, cells were treated with 100 nM Gemcitabine, and 16 hrs later were fixed and stained for the DNA damage markers γH2AX and 53BP1. Cells with >5 nuclear foci were scored as positive. All experiments were performed independently at least three times. Statistical analysis was conducted by one-way ANOVA. ns=not significant, *p<0.05, ***p<0.001, ****p<0.0001

To assess the impact of PRIMPOL on Gemcitabine resistance, we performed MTT-based survival assays. As before, MRNIP KO cells were markedly resistant to Gemcitabine relative to WT HeLa cells. PRIMPOL depletion had minimal effect on WT cell drug sensitivity, but partially re-sensitised MRNIP KO cells, consistent with the elevated DNA gaps observed under these conditions (Fig 3C). Similar findings were obtained following PRIMPOL depletion in WT and MRNIP KO HCT116 cells (Supplementary Fig 2D). Survival data obtained following UBC13 and REV3L depletion was not included, because depletion of either factor had profound negative effects on the long-term viability of both WT and MRNIP KO cells, even in the absence of genotoxic challenge. We then tested the effects of UBC13, PRIMPOL and REV3L depletion on the prevalence of DNA damage markers in Gemcitabine-treated WT and MRNIP KO cells. Gemcitabine treatment led to an increase in the proportion of WT cells positive for γH2AX foci. As we have reported previously, an increased proportion of unchallenged MRNIP KO cells exhibit γH2AX foci. However, Gemcitabine treatment did not induce an additional increase in γH2AX foci in MRNIP KO cells, which is consistent with the relative CTNA resistance of this line (Fig 3D). Furthermore, UBC13, PRIMPOL or REV3L depletion selectively led to a marked elevation in the proportion of γH2AX foci-positive MRNIP KO but not WT cells following Gemcitabine treatment (Fig 3E-H). The efficacy of PRIMPOL and UBC13 knockdowns were assessed by Western blotting, while REV3L depletion was tested via qRT-PCR, given the absence of a high-quality specific REV3L antibody (Fig 3E). Our findings support a model in which MRNIP loss limits DNA damage and chemosensitivity by facilitating enhanced Gemcitabine removal and subsequent post-replicative gap filling via template switch, driven by POL-ζ.

### Primase redundancy in Gemcitabine-treated MRNIP KO cells

That PRIMPOL depletion leads to restoration of DNA gaps in CTNA-treated MRNIP KOs initially appeared paradoxical, given the extensive published data demonstrating that PRIMPOL loss suppresses DNA gaps generated in response to a range of replication stress-inducing agents^6,7,20,24–27^. PRIMPOL exhibits both primase and polymerase activities. To test whether the observed DNA gaps in PRIMPOL-depleted MRNIP KO cells are associated with primase or polymerase functions, we generated MRNIP KO cells stably expressing either WT or CH mutant PRIMPOL, in which mutation of Cys419 and His426 disrupts primase activity while preserving polymerase activity^1^. To avoid siRNA targeting of ectopic PRIMPOL mRNA, we employed an siRNA targeting the PRIMPOL untranslated region (UTR). As observed previously, depletion of PRIMPOL resulted in restoration of DNA gaps in Gemcitabine-treated MRNIP KO cells, demonstrating that this phenotype is reproducible with two independent PRIMPOL siRNAs. Expression of WT but not CH mutant PRIMPOL in PRIMPOL-depleted MRNIP KO cells suppressed Gemcitabine-induced DNA gaps, suggesting that the observed phenotype is attributable to the primase function of PRIMPOL (Fig 4A). Expression of WT or CH mutant PRIMPOL in this system was validated by western blotting (Fig 4B).

**Figure 4:**
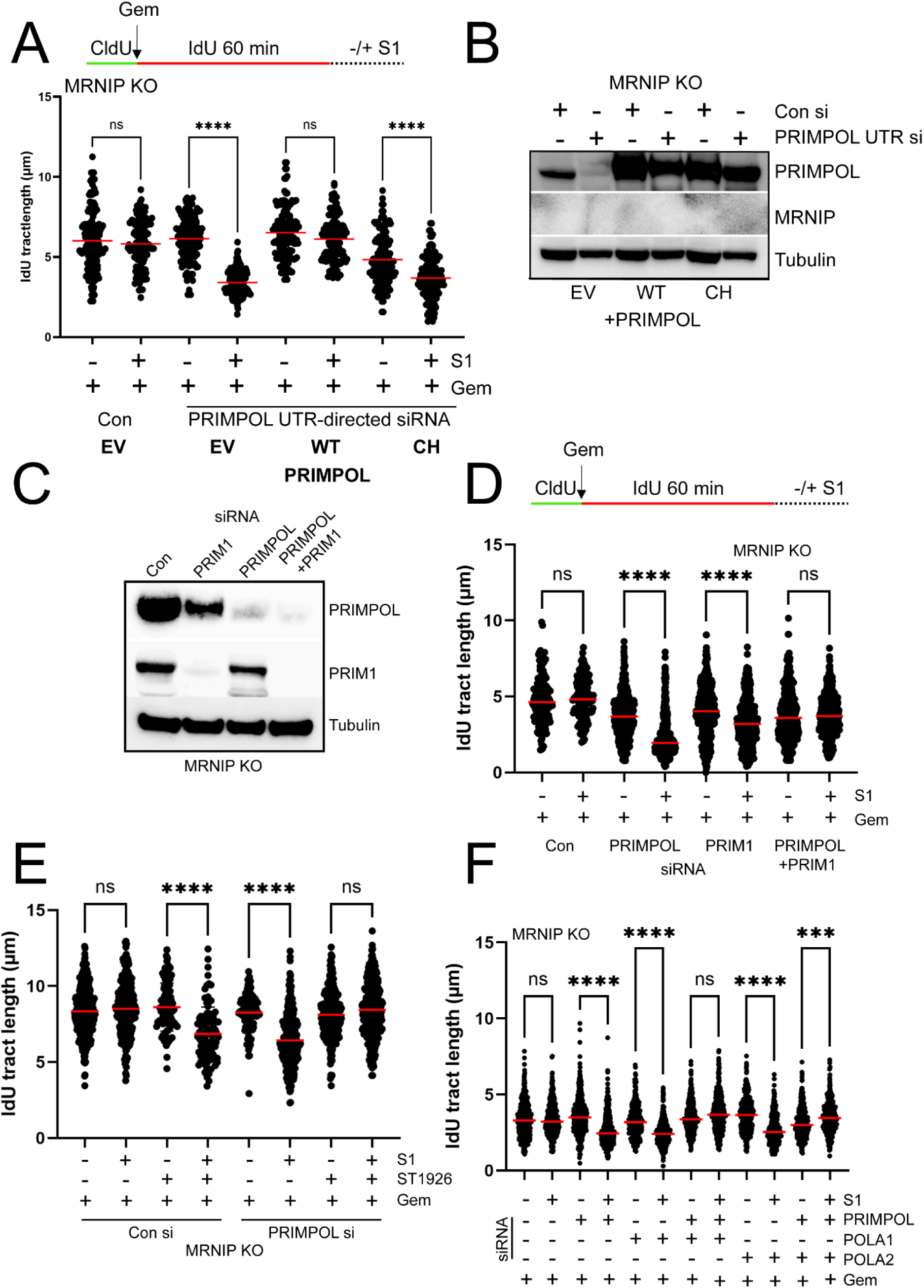
Primase redundancy in Gemcitabine-treated MRNIP KO cells. A and B: MRNIP KO cells stably expressing WT or CH mutant PRIMPOL were transfected with a control siRNA, or a UTR-directed PRIMPOL siRNA. After 48 hrs, cells were labelled with CldU followed by Gemcitabine and IdU, and DNA gaps were assessed by S1-linked DNA fibre assay (**A**). PRIMPOL status was monitored by western blotting (**B**)**. C and D:** MRNIP KO HeLa cells were transfected with a control siRNA, or siRNAs targeting PRIM1 or PRIMPOL alone and in combination. Depletion of both factors was monitored by western blotting (**C**). After 48 hrs, cells were treated as in (A) and DNA gaps assessed via S1-linked DNA fibre assay (**D**). **E:** MRNIP KO cells were transfected with a control siRNA or PRIMPOL siRNA, and after 48 hrs cells were pre-treated with DMSO or 1 μM ST1926 in combination with 100 nM Gemcitabine. S1-linked DNA fibre assays were performed as in (A). **F:** MRNIP KO HeLa cells were transfected with a control siRNA, or siRNAs targeting POLA1, POLA2 or PRIMPOL both alone and in combination. After 48 hrs, cells were treated as in (A) and DNA gaps assessed via S1-linked DNA fibre assay. All experiments were performed independently least three times. Statistical analysis was conducted by one-way ANOVA. ns=not significant, ***p<0.001, ****p<0.0001

The genome of higher eukaryotes encodes only two enzymes with recognised primase activity – PRIMPOL and the Pol α-primase complex, the latter of which is comprised of PRIM1, PRIM2, POLA1 and POLA2 subunits^29^. To test whether DNA primase is responsible for generation of DNA gaps in MRNIP KO cells in the absence of PRIMPOL, we co-depleted PRIMPOL and PRIM1, which is the catalytic subunit of DNA primase. PRIMPOL and PRIM1 depletion was validated by western blotting (Fig 4C). Strikingly, although depletion of either PRIMPOL or PRIM1 alone resulted in DNA gaps, co-depletion of both PRIMPOL and PRIM1 led to DSG suppression, suggestive of primase redundancy following chain termination (Fig 4D). We did not observe decreased fork progression in PRIM1-depleted cells, despite robust knockdown. It is possible that RNAi-mediated depletion selectively affects free primase complexes, and that minimal residual PRIM1 protein levels are sufficient to build a stable set of primase-proficient replisomes. We also note that prior publications report only moderate defects in replication fork progression in cells housing a PRIM1 mutation^28^. Given the novelty of this phenotype, we conducted additional experiments to test whether restoration of DNA gaps in PRIMPOL-depleted cells is reversed by perturbation of other Pol α-primase components. We treated PRIMPOL-depleted MRNIP KO cells with the Pol-α inhibitor ST1926. or depleted MRNIP KO cells of PRIMPOL in combination with depletion of POLA1 or POLA2. Whilst individual POLA1 or POLA2 depletion or ST1926 pre-treatment resulted in post-replicative DNA gaps as previously reported^30^, the same perturbations led to suppression of DNA gaps induced by PRIMPOL depletion (Fig 4E and F and Supplementary Fig 3A), providing further support for the hypothesis that DNA gaps in PRIMPOL-depleted MRNIP KO cells are dependent on the functionality of Pol α-primase.

### DNA gaps in PRIMPOL-depleted MRNIP KO cells are modulated by fork reversal, MRE11, 53BP1 and EXO1

Replication repriming suppresses replication fork reversal^25,31,32^. Therefore, deficiency in repriming on either the leading or lagging strands following genotoxic challenge with CTNAs is likely to lead to increased reversal. We hypothesised that following defective repriming, dysregulated MRE11 resects the 3′ end of the regressed arm of the reversed fork to generate an overhang that can serve as a template on which the remaining functional primase can act. To test this hypothesis, we performed S1-linked fibre assays in PRIMPOL-depleted MRNIP KO cells following perturbation of SMARCAL1 or MRE11. Depletion of SMARCAL1 in PRIMPOL-depleted MRNIP KO cells caused DSG suppression, suggesting that in the absence of PRIMPOL, primosome-dependent DNA gaps are formed downstream of fork reversal (Fig 5A and B). Treatment of PRIMPOL-depleted MRNIP KO cells with PFM39 or Mirin also resulted in DSG suppression, demonstrating that formation of primosome-driven DNA gaps is dependent on the MRE11 exonuclease (Fig 5C). This is consistent with the hypothesis that the exonucleolytic digestion of reversed fork DNA ends facilitates redundant repriming at chain termination sites.

**Figure 5:**
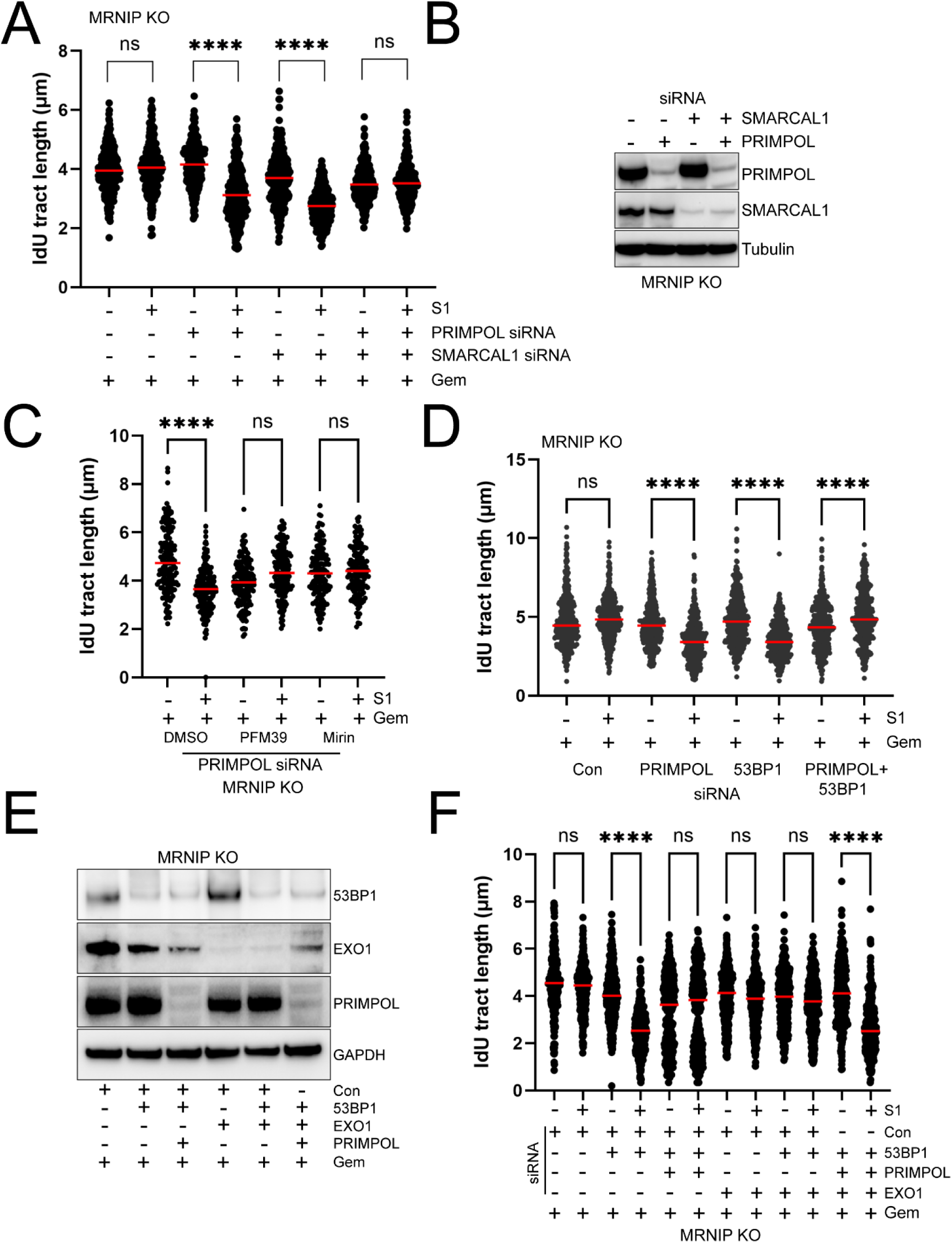
DNA gaps in PRIMPOL-depleted MRNIP KO cells are modulated by fork reversal, MRE11, 53BP1 and EXO1. A and B: MRNIP KO cells were transfected with a control siRNA, or siRNAs targeting PRIMPOL or SMARCAL1, both individually and in combination. After 48 hrs, cells were labelled with CldU followed by Gemcitabine and IdU, and DNA gaps were assessed by S1-linked DNA fibre assay (**A**). SMARCAL1 and PRIMPOL status was monitored by western blotting (**B**)**. C:** MRNIP KO HeLa cells were transfected with an siRNA targeting PRIMPOL. After 48 hrs, cells were treated with DMSO, or 25 μM PFM39 or Mirin, then S1-linked DNA fibre assays were conducted as in (A). **D:** MRNIP KO cells were transfected with a control siRNA, or siRNAs targeting PRIMPOL or 53BP1, alone and in combination. After 48 hrs, S1-linked DNA fibre assays were conducted as in (A). **E and F:** MRNIP KO cells were transfected with a control siRNA, or siRNAs targeting PRIMPOL, 53BP1 or EXO1, alone and in combination. After 48 hrs, S1-linked DNA fibre assays were conducted as in (A). All experiments were performed independently least three times. Statistical analysis was conducted by one-way ANOVA. ns=not significant, ****p<0.0001

The DSB repair pathway choice factor 53BP1 has been extensively implicated in modulation of end-resection and has been reported to protect a subset of reversed replication forks from nucleolytic degradation^33,34^. Studies have also suggested that 53BP1 acts to constrain the digestion of DNA ends by the 5’-3’ nuclease EXO1, which acts at DSBs, DNA gaps and reversed replication forks^35^. We performed S1-linked fibre assays in Gemcitabine-treated MRNIP KO cells following depletion of 53BP1 or PRIMPOL, both alone and in combination. This experiment revealed that depletion of 53BP1 results in DSG restoration in MRNIP KO cells, although DNA gaps observed in PRIMPOL-depleted cells were suppressed by co-depletion of 53BP1 (Fig 5D). Furthermore, depletion of EXO1 suppressed DNA gaps associated with 53BP1 depletion in PRIMPOL-proficient cells, while conversely EXO1 depletion led to restoration of DNA gaps suppressed by 53BP1 depletion in PRIMPOL-depleted cells (Fig 5E and F). This data is consistent with a model in which 53BP1 promotes gap filling by limiting the EXO1-dependent extension of post-replicative DNA gaps generated by PRIMPOL. In the absence of PRIMPOL, we hypothesise that 53BP1 facilitates primase redundancy at reversed forks by suppressing the EXO1-mediated digestion of the 5’ ends of regressed arm overhangs formed by limited MRE11-dependent digestion (see Model).

### CDK-dependent MRNIP phosphorylation is required for suppression of Gemcitabine-induced DNA gaps

Mass spectrometric analysis of FLAG-tagged MRNIP immune complexes revealed that MRNIP is phosphorylated on Ser217, a proline-directed site^36^. We raised a polyclonal antibody against a Ser217-containing phospho-peptide and tested the specificity of this antibody in FLAG antibody immune complexes derived from MRNIP KO HeLa cells stably expressing doxycycline-inducible FLAG-tagged WT or Ser217Ala MRNIP. This test confirmed that the antibody selectively recognises phosphorylated MRNIP (Figure 6A). MRNIP phosphorylation was markedly increased by overnight treatment with the microtubule depolymerising agent colcemid, which arrests cells in metaphase. Since Ser217 is a proline-directed site phosphorylated late in the cell cycle, we assessed Ser217 phosphorylation in presence and absence of RO-3306, an ATP-competitive inhibitor of CDK1. Again, S217 was markedly stimulated by colcemid treatment, and both basal and colcemid-stimulated MRNIP phosphorylation were reversed by RO-3306 pre-treatment, indicating that MRNIP phosphorylation is likely to be CDK1-dependent (Figure 5B). The hypothesis that Ser217 is a CDK-dependent site was further bolstered by our observation that Ser217 phosphorylation was significantly increased by pre-treatment with the WEE1 inhibitor Adavosertib (Fig 6C), which reduced CDK1 Y15 phosphorylation as anticipated. Adavosertib-induced MRNIP phosphorylation was abolished by co-treatment with RO-3306 (Fig 6C), demonstrating that the enhanced Ser217 phosphorylation associated with WEE1 inhibition is due to enhanced CDK1 activity.

**Figure 6:**
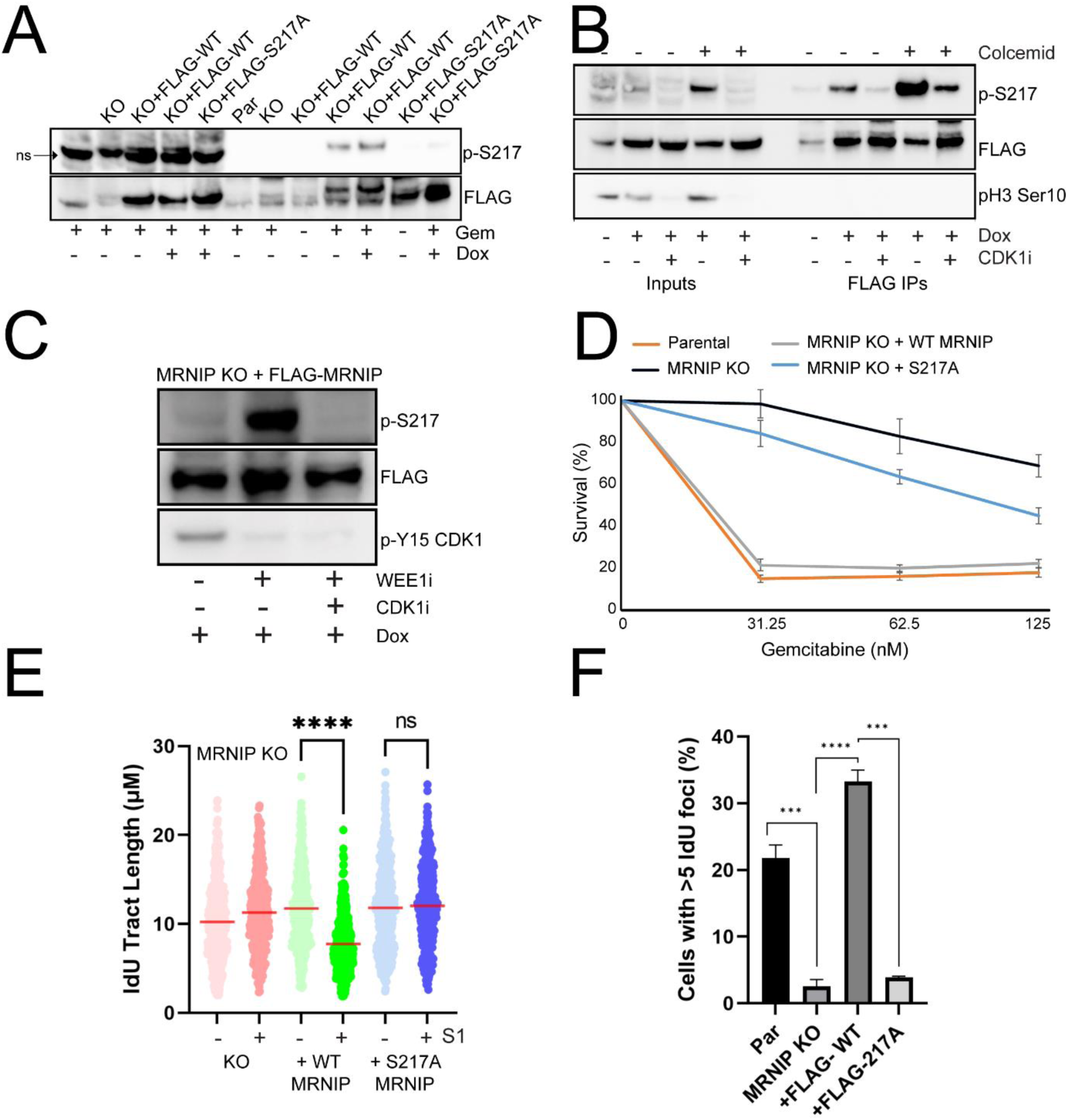
CDK-dependent MRNIP phosphorylation is required for DSG modulation. **A:** Parental HeLa cells, MRNIP KO cells or MRNIP KO cells stably expressing doxycycline-inducible FLAG-tagged WT or S217A MRNIP were treated with vehicle or 1 μg/ml doxycycline. After 24 hrs, cells were treated with vehicle control or 100 nM Gemcitabine for 16 hrs. Cells were then lysed, and FLAG-MRNIP enriched via immunoprecipitation with FLAG M2 antibody-coated beads. Immune complexes were eluted with 3xFLAG peptide, and eluates resolved by SDS-PAGE followed by western blotting with the indicated antibodies. **B:** MRNIP KO cells stably expressing doxycycline-inducible FLAG-tagged WT were treated with vehicle or 1 μg/ml doxycycline. After 16 hrs cells were treated with 100 ng/ml colcemid, then treated with either DMSO or 1μM RO-3306 for an additional 6 hrs. FLAG-MRNIP was immunoprecipitated, and FLAG immune complexes resolved by SDS-PAGE followed by western blotting with the indicated antibodies. **C**: MRNIP KO cells expressing FLAG-MRNIP were treated with 1 μM Adavosertib or 1 μM RO-3306, alone or in combination. After 4 hrs, cells were lysed and extracts probed with the indicated antibodies. **D:** MRNIP KO cells or MRNIP KO cells stably expressing doxycycline-inducible FLAG-tagged WT or S217A MRNIP were treated with vehicle or 1 μg/ml doxycycline. After 24 hrs, cells were treated with the indicated concentrations of Gemcitabine. After a further 72 hrs, cell viability was estimated by MTT assay. **E:** MRNIP KO cells or MRNIP KO cells stably expressing doxycycline-inducible FLAG-tagged WT or S217A MRNIP were treated with vehicle or 1 μg/ml doxycycline. After 24 hrs, cells were labelled with CldU then IdU in the presence of Gemcitabine, followed by mock digestion or ssDNA digestion with S1 nuclease. IdU tract length in the presence and absence of S1 was used as a readout of post-replicative ssDNA. **F:** MRNIP KO cells or MRNIP KO cells stably expressing doxycycline-inducible FLAG-tagged WT or S217A MRNIP were treated with vehicle or 1 μg/ml doxycycline. After 24 hrs, cells were labelled for 24 hrs with IdU, then treated with 100 nM Gemcitabine. After 16 hrs, cells were fixed and stained with an IdU cross-reactive BrdU antibody, and the proportion of cells with >5 nuclear foci was determined (F). Statistical analysis was conducted by one-way ANOVA. ns=not significant, **p<0.01, ***p<0.001, ****p<0.0001

To test whether Ser217 phosphorylation contributes to regulation of Gemcitabine sensitivity, we performed MTT-based viability assays in WT and MRNIP KO cells, and in MRNIP KO cells stably expressing FLAG-tagged WT or Ser217Ala MRNIP. We repeatedly attempted to generate cell lines stably expressing a FLAG-Ser217Asp phospho-mimetic MRNIP mutant but failed to recover colonies on all occasions. This suggests that mimicking constitutive phosphorylation of Ser217 negatively impacts cell growth or viability even in the relatively small quantities expressed in the absence of doxycycline, although this requires further investigation. As before, MRNIP KO cells were resistant to Gemcitabine, and expression of FLAG-WT MRNIP in the MRNIP KO background restored Gemcitabine sensitivity to WT levels (Figure 6D). In contrast, expression of the alanine substitution mutant of Ser217 failed to rescue Gemcitabine sensitivity, indicative of the functional relevance of this site (Figure 6D). Together, our data indicates that Ser217 is crucial for the functionality of MRNIP as a regulator of Gemcitabine resistance.

Next, we assessed the prevalence of DNA gaps and ssDNA in WT and Ser217Ala-expressing MRNIP KO cells via S1-linked fibre assays and native IdU, respectively. As previously observed, Gemcitabine-induced DNA gaps were suppressed in MRNIP KO cells. Stable expression of FLAG-WT MRNIP resulted in restoration of post-replicative ssDNA gaps, demonstrating that FLAG-tagged MRNIP remains functional in the context of gap regulation, and that ssDNA gap suppression in MRNIP KO cells is not an off-target effect of the CRISPR protocol. Notably, the Ser217Ala mutant did not modulate ssDNA gap prevalence in either assay, suggesting that phosphorylation of this site is an absolute requirement for MRNIP functionality in gap modulation (Fig 6E and F). Expression of FLAG-tagged proteins was confirmed via western blotting (Supplementary Fig 3B). Immunoprecipitation experiments with FLAG-tagged WT Ser217Ala MRNIP revealed that this site is not required for the interaction between MRNIP and MRE11 (Supplementary Fig 3C). Collectively, our data suggests a role for CDK1-mediated MRNIP phosphorylation in ssDNA gap suppression and Gemcitabine chemosensitivity, and a potential novel role for S-phase CDK activity in the regulation of post-replicative ssDNA gaps.

## Discussion

Treatment of cells with a wide range of replication stress inducers leads to the formation of repriming-dependent post-replicative DNA gaps, which must be filled in a timely manner to limit genome instability. The tumour suppressors BRCA1 and BRCA2 have been extensively linked with DSG suppression, and several studies support a mechanistic link between DSG prevalence and chemosensitivity, largely in the context of treatment with the cross-linking chemotherapy Cisplatin or PARP inhibition^6–13,27^. The broader regulatory mechanisms at play in DSG regulation remain largely uncharacterised.

CTNAs represent a subclass of nucleoside analogues that stall DNA replication via direct incorporation into the nascent DNA strand. The continuity of replication under these circumstances is dependent on CTNA removal or replication repriming. Given our prior discovery that the MRE11 regulator MRNIP modulates DNA gaps in response to Cisplatin and Olaparib and the recent finding that MRE11 can act to remove Gemcitabine from genomic DNA, we examined the regulation of DNA gaps formed by CTNAs, and the impact of MRNIP loss on DSG prevalence and chemoresistance. Our work reveals that MRNIP loss results in suppression of CTNA-induced DNA gaps and DNA damage and leads to chemoresistance. Our data suggests a mechanism via which MRE11-dependent activity drives the resection of chain-terminated DNA ends prior to UBC13 and REV3L-dependent post-replicative gap filling.

In contrast to treatment with other genotoxins that do not specifically target the nascent strand, we observe marked PRIMPOL-dependent post-replicative DNA gaps in WT cells treated with Gemcitabine and the obligate chain terminator Cytarabine. This suggests that repriming plays a primary role at sites of CTNA incorporation, which is perhaps to be expected given the direct block to nascent chain extension, at least in the absence of very rapid analogue removal. The persistence of DNA gaps under these conditions suggests that the kinetics of CTNA removal and/or the completion of gap-filling mechanisms are relatively inefficient.

In both MRNIP KO cells and cells depleted of MRNIP via RNAi, we observed suppression of CTNA-induced DNA gaps, which was surprisingly reversed by inhibition of the MRE11 exonuclease. At first this appeared contradictory, given the extensive evidence implicating MRE11 in DSG extension in multiple models^9,13^. However, we reasoned that in the absence of MRNIP, dysregulated MRE11 may more efficiently resect the chain-terminated 3’ end, leading not only to Gemcitabine removal but to additional resection that may be a requisite for the generation of a ssDNA stretch of sufficient length to license template switch-mediated gap filling. Indeed, depletion of the known template-switch/gap filling factors UBC13 or REV3L led to restoration of DSG in Gemcitabine-treated MRNIP KO cells. This data is consistent with work demonstrating that modulation of DNA resection factors dictates gap-filling pathway choice^37^.

More recent studies indicating the occurrence of bidirectional DSG resection and suggest a limit to the length of DNA gaps that can be successfully repaired^38,39^. Work from the Vindigni laboratory demonstrates that in BRCA1-deficient cells, DNA gaps can be more efficiently filled following the limitation of either 5’ or 3’ end resection via modulation of EXO1/DNA2/BLM or MRE11, respectively^39^. This adds weight to the hypothesis that the relatively poor processivity of TLS polymerases imposes a size limit on effective and timely gap filling. We examined the effect of depletion of the anti-resection factor 53BP1 on DSG dynamics in MRNIP-deficient cells. 53BP1 has been directly implicated in the control of EXO1-dependent resection^35^. Our work demonstrates that 53BP1 is required for suppression of DNA gaps in CTNA-treated MRNIP KO cells, and that gaps formed in the absence of 53BP1 are restored by co-depletion of EXO1. Since MRE11-dependent resection underpins gap filling in the absence of MRNIP, our data supports a model in which loss of 53BP1 and MRNIP leads to bidirectional loss of control of resection of post-replicative DNA gaps, supportive of the hypothesis that DNA gaps can become too wide to be effectively filled (see Model 1).

Whilst we demonstrated that CTNA-induced DNA gaps are a consequence of PRIMPOL activity in WT cells, we also consistently observed restoration of DNA gaps in CTNA-treated MRNIP KO cells following PRIMPOL depletion. Given that the human genome encodes only two known primases, we hypothesised that in the absence of PRIMPOL-dependent repriming, a secondary repriming mechanism takes place at nucleoside analogue-blocked ends. Indeed, depletion of the catalytic primosome components PRIM1, POLA1 and POLA2 or Pol α inhibition resulted in restoration of post-replicative gaps, while co-depletion of primosome factors and PRIMPOL resulted in gap suppression. These observations suggest the existence of primase redundancy, at least in the context of CTNA-mediated nascent strand blockade. Pol α has been demonstrated to catalyse replication on leading strand forked structures *in vitro*^40^, and the action of Pol α-primase outside the context of the lagging strand has been proven by other studies^41^. There has been speculation that PRIMPOL evolved in higher eukaryotes at least in part as a means of compensation for the inefficient leading strand repriming mediated by Pol α^41^. Impairment of fork reversal via SMARCAL1 depletion suppressed DNA gaps in PRIMPOL-depleted MRNIP KO cells, which is consistent with a model in which repriming suppresses fork reversal. Since MRE11 exonuclease inhibition suppressed gaps associated with PRIMPOL depletion, we suggest that gap formation in the absence of repriming is dependent on resection of reversed fork DNA. Since primosome-mediated repriming requires a suitable template, we hypothesised that MRE11-dependent 3′ end resection of regressed nascent DNA at reversed forks creates a 5′ overhang on which Pol α-primase can reprime replication in the absence of PRIMPOL, and which represents a secondary mechanism of DSG generation. In support of this hypothesis, we found that 53BP1 depletion suppressed DNA gaps in PRIMPOL-depleted cells, and that gaps were restored by the additional depletion of EXO1. These findings support a model in which 53BP1 limits EXO1-dependent 5’ end resection at reversed forks to maintain a template overhang that facilitates primosome-dependent repriming. This effect has not been widely observed in PRIMPOL-deficient cells in response to other genotoxins and may be specific to scenarios involving CTNA-blocked sites – this requires further investigation beyond the scope of this manuscript. We speculate that our findings represent a novel redundancy mechanism via which defective repriming on either the lagging (PRIM1-mediated) or leading (PRIMPOL-mediated) strands can be resolved via fork remodelling.

The context-specific activity of 53BP1 in our experiments is interesting. 53BP1 is recruited to DSBs via binding of the Tudor domain to modified Histones H2 and H4, although a recent report suggests that 53BP1 directly binds PRIM1-dependent RNA species at replication forks^42^. Furthermore, recent work from the Lambert laboratory demonstrates the existence of protective KU70/80-bound primase-dependent RNA at reversed forks^43^, although whether this RNA is integrated into the leading or lagging strands was not elucidated. It is possible that 53BP1 limits EXO1-driven resection of PRIMPOL-dependent post-replicative DNA gaps via canonical binding of the Tudor domain to DSG-proximal H2K15Ub and H4K20me2 at post-replicative ssDNA-dsDNA junctions. Alternatively, given that nucleosomes are dismantled at replication forks to facilitate replisome progression, 53BP1 might be recruited to RNA primers formed at reversed forks in the absence of canonical PRIMPOL-dependent repriming.

During this study, we also identified the proline-directed phosphorylation of Ser217 as crucial to MRNIP function in DSG regulation. Experiments with the inhibitor RO-3306 suggested the involvement of CDK1 in mediating phosphorylation of this site, suggestive of a broader role for CDK1 in DSG regulation. Indeed, pre-treatment with RO-3306 also prevented Gemcitabine-induced DNA gaps in WT HeLa cells (data not shown), although the role of CDK1 in this context requires further investigation. ATR inhibition has been linked to CDK1-dependent dormant origin firing^44^, and we noted delayed CHK1 activation following Gemcitabine treatment (data not shown). We attempted to employ analogue-sensitive CDK1-AS HeLa cells, which are modified to allow the specific inhibition of CDK1 following treatment with the bulky analogue 1NM-PP1. Although we observed robust suppression of DNA gaps in 1NM-PP1-treated CDK1-AS cells, subsequent experiments demonstrated that analogue treatment suppressed CTNA-induced DNA gaps even in unmodified HeLa cells, demonstrating off-target effects of this compound (data not shown). Exactly why Ser217 is important is currently unclear.

Collectively, our findings demonstrate that nuclease dysregulation can license post-replicative gap filling mechanisms and drive chemoresistance and add mechanistic insight into replication dynamics following CTNA challenge. Whilst our work may have broader implications in cancer treatment, further research is required to investigate the possibility that loss of nuclease regulation in tumours constitutes an innate or acquired mechanism of therapeutic resistance.

## Study Limitations

Our work has been performed exclusively in a two-dimensional cell culture setting. Further work is required to elucidate the *in vivo* roles of MRNIP, and to test whether MRNIP-dependent gap modulation in this context is of deeper physiological relevance.

## Acknowledgements

We thank Prof Juan Mendez for the kind gift of PRIMPOL antibody.

## Author contributions

CJS, DH, LGB, VT and EV. devised and performed the majority of experiments including all CRISPR line generation and validation, and DNA fibre assays with associated analysis and data formatting. CJS wrote the paper with contributions from EH and assisted with immunofluorescence and survival assays. VT, DH and JR performed native IdU assays. EV, EH and RK assisted with mass spectrometric analysis. FA contributed to survival assays and some IF experiments. CJ contributed survival data.

## Data and materials availability

All data needed to evaluate the conclusions in the paper are present in the paper. Additional data related to this paper may be requested from the authors.

## Model 1.

LHS: Gemcitabine incorporation leads to replication fork stalling and PRIMPOL-dependent replication repriming to generate a nascent strand ssDNA discontinuity. In WT cells, phosphorylated MRNIP limits MRE11-dependent 3’ end resection of the ssDNA gap. In MRNIP KO cells, enhanced DSG resection by MRE11 leads to Gemcitabine removal and UBC13/REV3L-dependent gap filling by template switch. 53BP1 limits EXO1 activity to prevent excessive bidirectional resection RHS: In the absence of PRIMPOL, Pol α-primase mediates repriming at MRE11-resected reversed forks, which is contingent upon template maintenance by 53BP1-mediated restraint of EXO1-driven 5’ end resection.

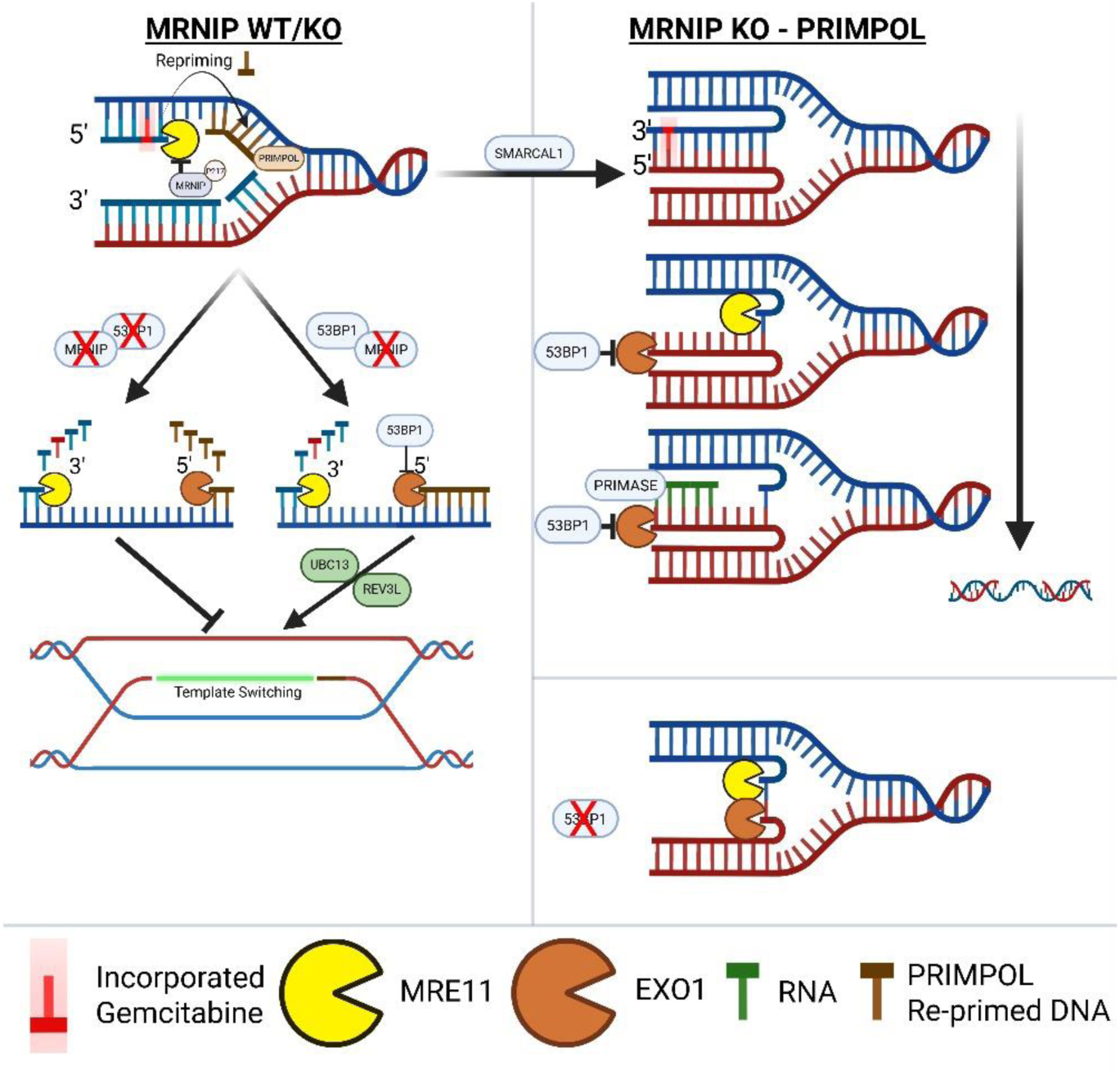

---

**Supplementary Figure 1.**
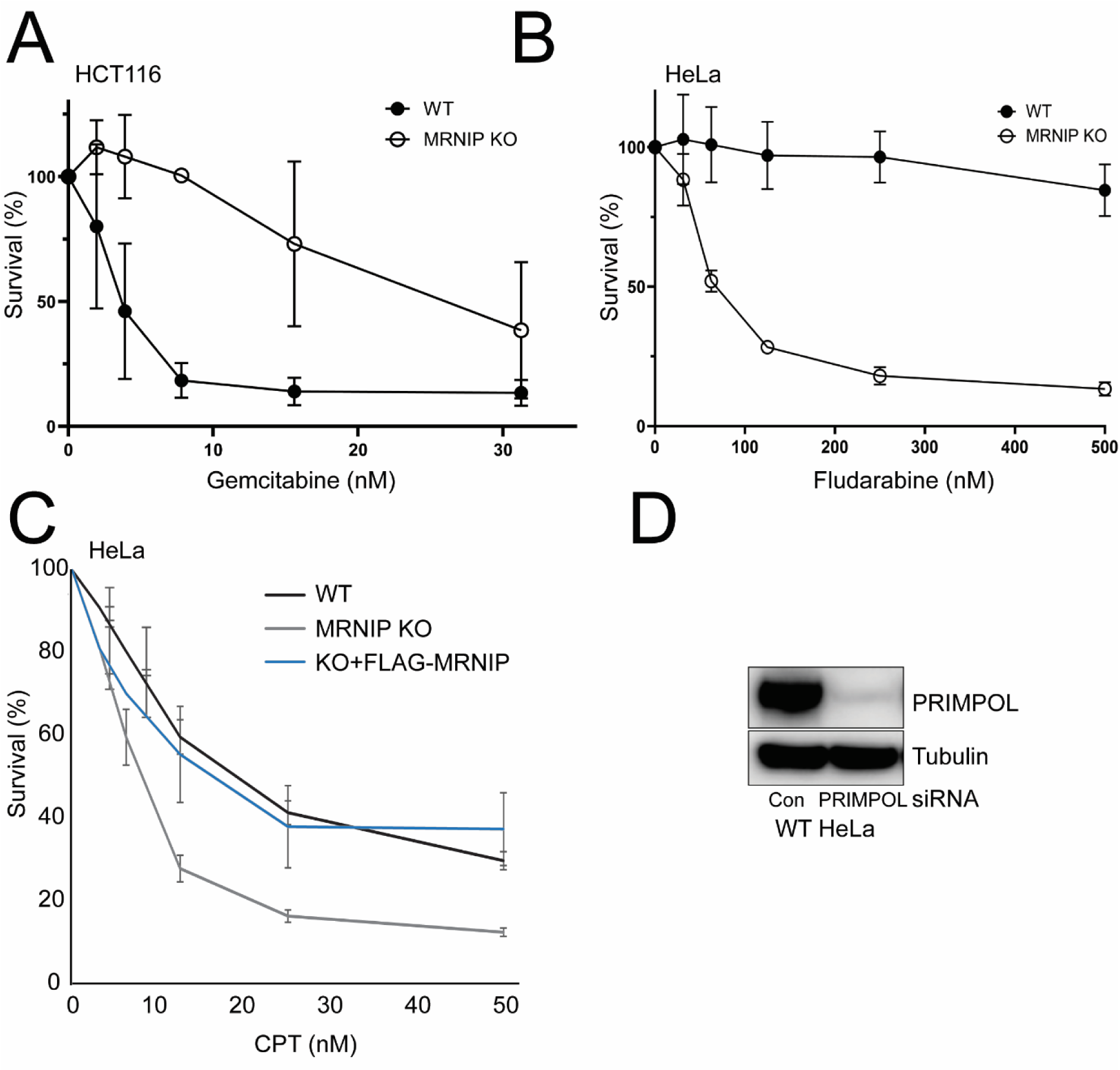
**A-C:** WT and MRNIP KO HCT116 or HeLa cells and MRNIP KO HeLa cells stably expressing FLAG-MRNIP were treated with the indicated concentrations of Gemcitabine, CPT or Fludarabine as indicated. After 72 hrs, cell viability was estimated by MTT assay, and data normalised to untreated controls. **D:** WT HeLa cells were transfected with a control siRNA, or an siRNA targeting PRIMPOL. After 48 hrs, cells were lysed and Western blotting employed to confirm PRIMPOL depletion. All experiments were performed at least three times, independently. Statistical analysis was conducted by one-way ANOVA. ns=not significant, ****p<0.0001

**Supplementary Figure 2.**
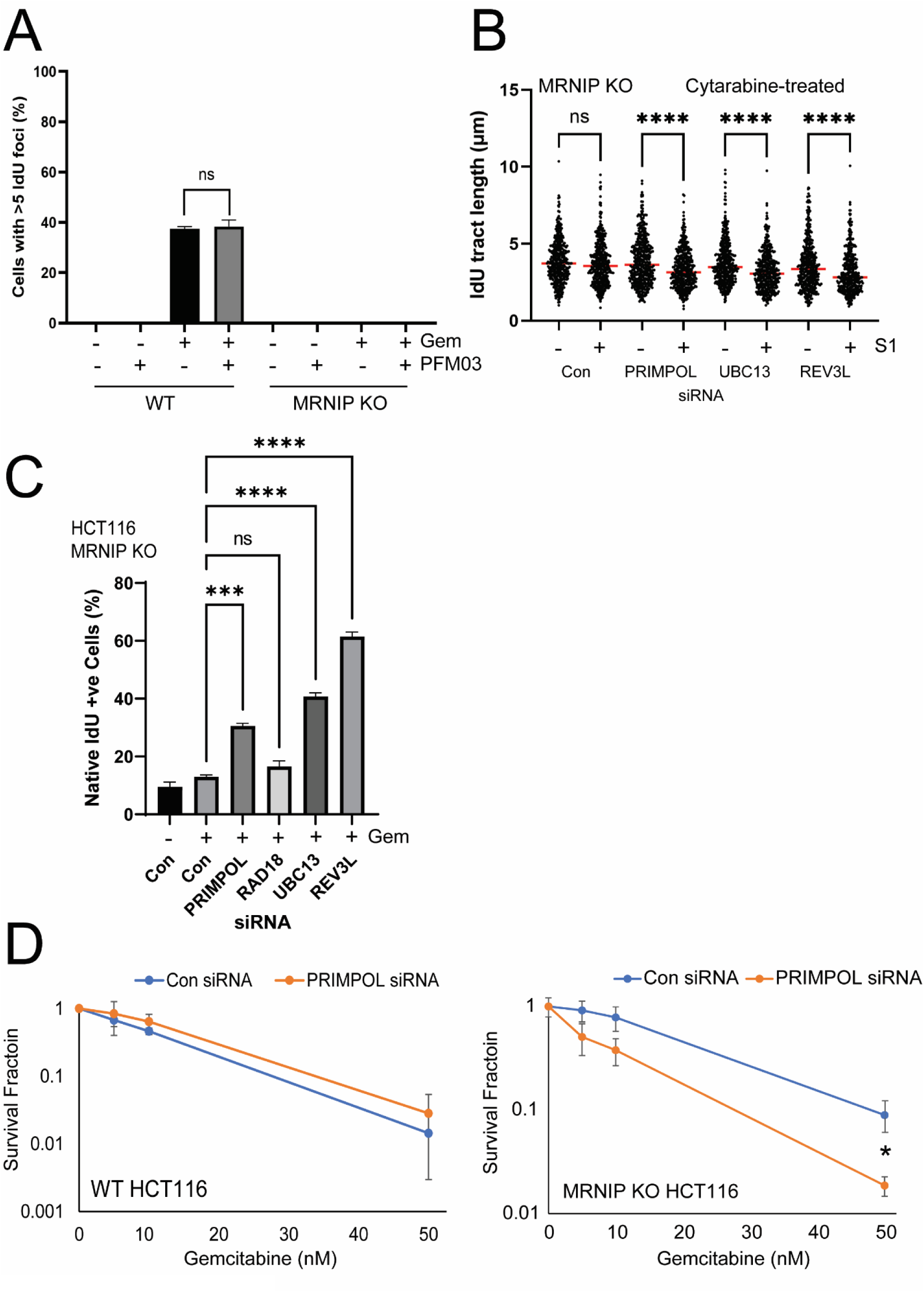
**A:** WT and MRNIP KO HeLa cells were labelled with IdU for 16 hrs, followed by 16 hrs treatment with vehicle control or 100 nM Gemcitabine in the presence of DMSO or 25 μM PFM03. Cells were then fixed and stained under native conditions with a IdU cross-reactive BrdU antibody, and counterstained with DAPI. Cells were assessed by confocal microscopy, and the proportion of cells with >5 nuclear IdU foci was determined. **B:** MRNIP KO cells were transfected with the indicated siRNAs. After 48 hrs, nascent DNA was CldU-labelled for 20 mins, followed by IdU labelling for 1 hr in the presence of 100 nM Cytarabine. ssDNA was then digested via S1 nuclease treatment, and DNA fibre spreading was performed. **C**: MRNIP KO HCT116 cells were transfected with the indicated siRNAs and labelled with IdU for 16 hrs, followed by 16 hrs treatment with 100 nM Gemcitabine. Cells were then fixed and stained under native conditions with a IdU cross-reactive BrdU antibody, and counterstained with DAPI. Cells were assessed by confocal microscopy, and the proportion of cells with >5 nuclear IdU foci was determined. **D:** WT and MRNIP KO HCT116 cells were transfected with a control siRNA or an siRNA targeting PRIMPOL. After 48 hrs cells were replated and a clonogenic survival assay performed in the presence of the indicated concentrations of Gemcitabine. Colony counts were normalised to untreated controls. All experiments were performed three times, independently. *p<0.05, ***p<0.001, ****p<0.0001

**Supplementary Figure 3.**
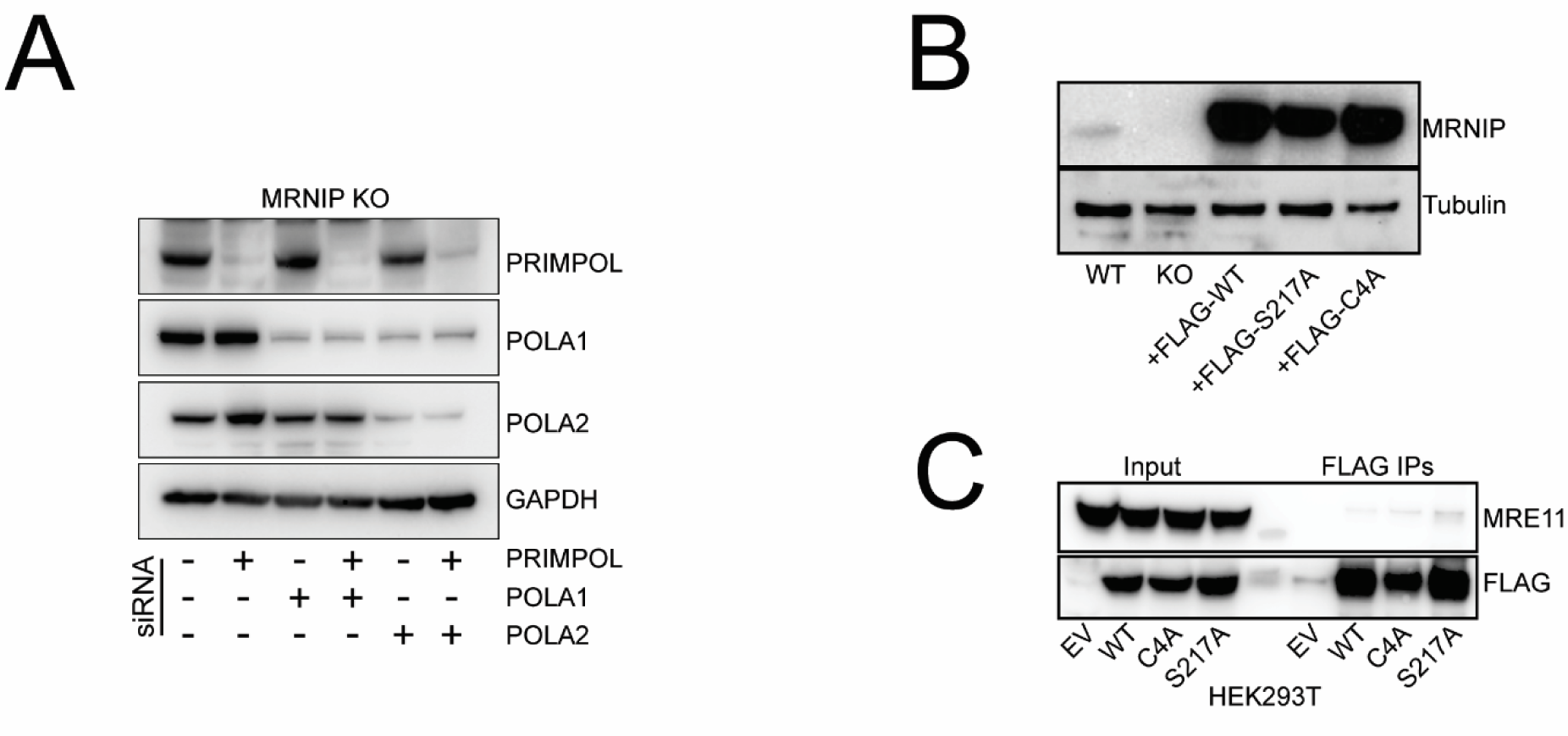
**A:** HeLa cells were transfected with a control siRNA, or siRNAs targeting PRIMPOL, POLA1 or POLA2 alone or in combination. After 48 hrs cells were lysed, and depletion confirmed by Western blotting with the indicated antibodies. **B:** HeLa cells stably expressing FLAG-tagged WT or Ser217Ala MRNIP were lysed, and MRNIP status was monitored by western blotting (G). Expression of the Cys4Ala mutant was also tested but not assessed. **C:** Cells as used in (A) were lysed and FLAG M2 antibody was used to purify WT and S217A MRNIP. Immune complexes were resolved by SDS-PAGE and probed with the indicated antibodies.

